# Optimizing immunostaining of archival fish samples to enhance museum collection potential

**DOI:** 10.1101/2022.07.21.501016

**Authors:** Garfield T. Kwan, Benjamin W. Frable, Andrew R. Thompson, Martin Tresguerres

**Affiliations:** Marine Biology Research Division, Scripps Institution of Oceanography, University of California San Diego, USA; Marine Vertebrate Collection, Scripps Institution of Oceanography, University of California San Diego, USA; NOAA Fisheries Service, Southwest Fisheries Science Center, La Jolla, CA, USA

**Keywords:** Quenching, antigen-retrieval, formalin, formaldehyde, immunohistochemistry, sodium borohydride

## Abstract

Immunohistochemistry (IHC) is a powerful biochemical technique that uses antibodies to specifically label and visualize proteins of interests within biological samples. However, fluid-preserved specimens within natural history collection often use fixatives and protocols that induce high background signal (autofluorescence), which hampers IHC as it produces low signal-to-noise ratio. Here, we explored techniques to reduce autofluorescence using sodium borohydride (SBH), citrate buffer, and their combination on fish tissue preserved with paraformaldehyde, formalin, ethanol, and glutaraldehyde. We found SBH was the most effective quenching technique, and applied this pretreatment to the gill or skin of 10 different archival fishes – including specimens that had been preserved in formalin or ethanol for up to 65 and 37 years, respectively. The enzyme Na^+^/K^+^-ATPase (NKA) was successfully immunostained and imaged using confocal fluorescence microscopy, allowing for the identification and characterization of NKA-rich ionocytes essential for fish ionic and acid-base homeostasis. Altogether, our SBH-based method facilitates the use of IHC on archival samples, and unlocks the historical record on fish biological responses to environmental factors (such as climate change) using specimens from natural history collections that were preserved decades to centuries ago.

**Highlights:**

- Sodium borohydride pretreatment reduced aldehyde-induced autofluorescence
- Successfully immunostained archival samples of various fixative and fixation time
- Larval fish that was formalin-fixed for 63-65 years was successfully immunostained

## Introduction

The potential of formalin as a histological preservation agent was first recognized in the late 19^th^ century (Simmons, 2014). By the early 20^th^ century, formalin was widely used in natural history collections to indefinitely preserve biological materials for morphological and taxonomic research (Simmons, 2014). Today, natural history collections contain an unparalleled array of biological specimens collected over the past centuries. As a result, natural history collections are often regarded as ‘time machines’ that allow researchers to examine and analyze samples collected decades to centuries ago (Shaffer et al., 1998). However, long-term formalin preservation is known to denature DNA (Stollar and Grossman, 1962), alter δ13C and δ15N isotope signatures (Edwards et al., 2002; Kelly et al., 2006), and induce protein crosslinkage that impedes immunohistochemistry (IHC; Pikkarainen et al., 2010; Stradleigh and Ishida, 2015). Past and ongoing research are aimed at improving DNA extraction (reviewed in Card et al., 2021; Raxworthy and Smith, 2021) and isotopic analysis (Hetherington et al., 2019; reviewed in Sarakinos et al., 2002) from archival specimens, whereas this study focuses on optimizing IHC of archival fish samples to unlock information about protein abundance and localization.

IHC is a biochemical technique that uses antibody-antigen immunoreactivity to label proteins of interest. Although tissues can be fixed with alcohol- or aldehyde-based fixatives, paraformaldehyde (PFA) is often preferable as it provides more optimal signal-to-noise ratio during imaging (Clancy and Cauller, 1998; Matsuda et al., 2011; Diez-Fraile et al., 2012). However, PFA is significantly more expensive than formalin, and likely will not be widely adopted as the preservation agent across biological archives. Moreover, many of the fluid-preserved specimens within natural history collections around the world have been fixed with or remained immerse within formalin. Thus, there is a great benefit to develop IHC techniques applicable to formalin-fixed samples stored within natural history collections.

Formalin fixation adversely affects IHC by inducing excessive protein crosslinkage, which prevents proper antibody binding (c.f. Pikkarainen et al., 2010; Stradleigh and Ishida, 2015). Furthermore, formalin fixation results in high levels of background signal, which masks the antibody-dependent signal and results in low signal-to-noise ratio (reviewed in Shi et al., 2011; Stradleigh and Ishida, 2015). One strategy often used to improve signal-to-noise ratio is by “quenching” background autofluorescence and endogenous peroxidase activity (reviewed in Shi et al., 1997, 2011). These quenching protocols are typically optimized for paraffin-embedded mammalian tissue sections using reagents such as sodium borohydride (SBH) or citrate buffer (CB) (Shi et al., 1993, 2011; Baschong et al., 2001; Luquin et al., 2010; Oliveira et al., 2010; Matsuda et al., 2011). In contrast, research dedicated to developing quenching protocols for formalin-fixed, non-mammalian archival tissue has been scant. In fact, IHC studies attempting to revive archival fish tissue for whole-mount imaging has been virtually nonexistent. Despite this lack of research, several studies have successfully used fluorescent IHC to immunotarget the enzyme Na^+^-K^+^-ATPase (NKA) in formalin-fixed fish gill (Wilson et al., 2000; Kwan et al., 2019; Frommel et al., 2021), pseudobranch (Yang et al., 2014), and intestine (Esbaugh and Cutler, 2016). However, these successful immunostaining on formalin-fixed samples required targeted dissection (only the tissue of interest was fixed), transfer of samples from fixatives to alcohol within hours (typically 8 – 24 hrs), quenching procedures, sectioning, and/or the use of confocal microscopy. Importantly, the targeted sampling and fixation duration sharply contrast with existing collection protocols: curators and collection managers are tasked with fixing an intact fish, which depending on its size, may take weeks to months for formalin to permeate through the entire specimen. Furthermore, there may be limited reagents available during research collections, especially when at sea or remote field stations. As a result, the fixation duration of archival samples varies greatly, with some samples being preserved from weeks to years at a time. Given the immense amount of historical data chronicled within archival samples, it is necessary to test, develop, and validate a robust quenching protocol that is applicable to natural history collections.

Natural history collections also contain specimens fixed in other preservatives. Ethanol is one of the oldest preservatives, and it is believed to have been used since the 13^th^ century (Simmons, 2014). Because ethanol was relatively expensive and distorts specimens, its popularity (as a fixative) declined as the relatively cheaper formalin became the dominant preservation agent in the 20^th^ century. More recently, ethanol has renewed utilization since samples fixed within >95% ethanol can preserve DNA (Card et al., 2021; Raxworthy and Smith, 2021) and do not dissolve CaCO_3_ microstructures such as otoliths (Swalethorp et al., 2016). Next, glutaraldehyde (GTA) is another preservative sometimes used in museums that fixes tissue more effectively, but at a slower rate than formaldehyde. Thus, GTA is sometimes used in combination with formaldehyde to enhance strength and rate of fixation (Simmons, 2014; Stradleigh and Ishida, 2015), and their excellent preservation of cell ultrastructure facilitates scanning electron microscopy (SEM). Even though samples fixed with 95% ethanol or GTA are not intended for IHC (Kiernan, 2000; Matsuda et al., 2011), they may be viable alternatives if the background signal could be sufficiently dampened, and their immunoreactivity recovered.

This study aims to identify and validate a robust IHC technique on archived fish samples preserved with fixatives at durations that reflect those commonly found within natural history collections. In this study, wild-caught splitnose rockfish (*Sebastes diploproa*) gills were fixed in 4% PFA, 10% formalin, 20% formalin, 95% ethanol, and combined 3% PFA, 0.35% GTA in 0.1M sodium cacodylate buffer (SEM fixative) at durations that reflect both experimental and archival sampling. Next, we tested the effectiveness of SBH, CB, or combined SBH and CB quenching on rockfish gills, and quantified their viability on reducing autofluorescence. Next, we validated the most promising of the three quenching treatments on archival samples housed at the *Scripps Institution of Oceanography (SIO) Marine Vertebrate Collection* (La Jolla, CA, USA) and the *CalCOFI Ichthyoplankton Collection* (La Jolla; La Jolla, CA, USA), some of which were preserved in formalin and ethanol for as long as 65 and 37 years, respectively (Table 1). The *SIO Marine Vertebrate Collection* contains samples dating back to 1884, and houses ~2,000,000 fishes across ~5,700 species (Singer et al., 2018; Frable, 2022). The majority of these samples are fixed in formalin, with the rest mainly preserved in ethanol. The *Ichthyoplankton Collection* at NOAA Southwest Fisheries Science Center (SWFSC) houses larval fish samples that have been consistently collected in the California Current Ecosystem by the California Cooperative Oceanic Fisheries Investigations (CalCOFI) since 1949, and includes ~75,000 larvae spanning across ~550 species (Moser et al., 2002). The majority of these specimens are also preserved in formalin, but since 1997, CalCOFI has preserved samples from the port side of bongo nets in 95% ethanol (Thompson et al., 2017). In addition, PFA- and GTA-fixed specimens preserved for experimental analysis were provided by the Tresguerres lab (SIO) and other research groups to be used as a comparison with archival samples. Fish samples were immunostained against the enzyme NKA due to the many existing studies that have successfully imaged NKA-rich ionocytes within the skin (c.f. Varsamos et al., 2002a; Kwan et al., 2019) and gill (c.f. Christensen et al., 2012; Montgomery et al., 2022) of various teleost fishes, and its essential role in maintaining ionic and acidbase homeostasis (reviewed in Varsamos et al., 2005; Evans et al., 2005; Glover et al., 2013). Altogether, 10 fish species were whole-mount immunostained against NKA and visualized with fluorescent secondary antibodies on epifluorescence and confocal microscopy.

**Table 1.**
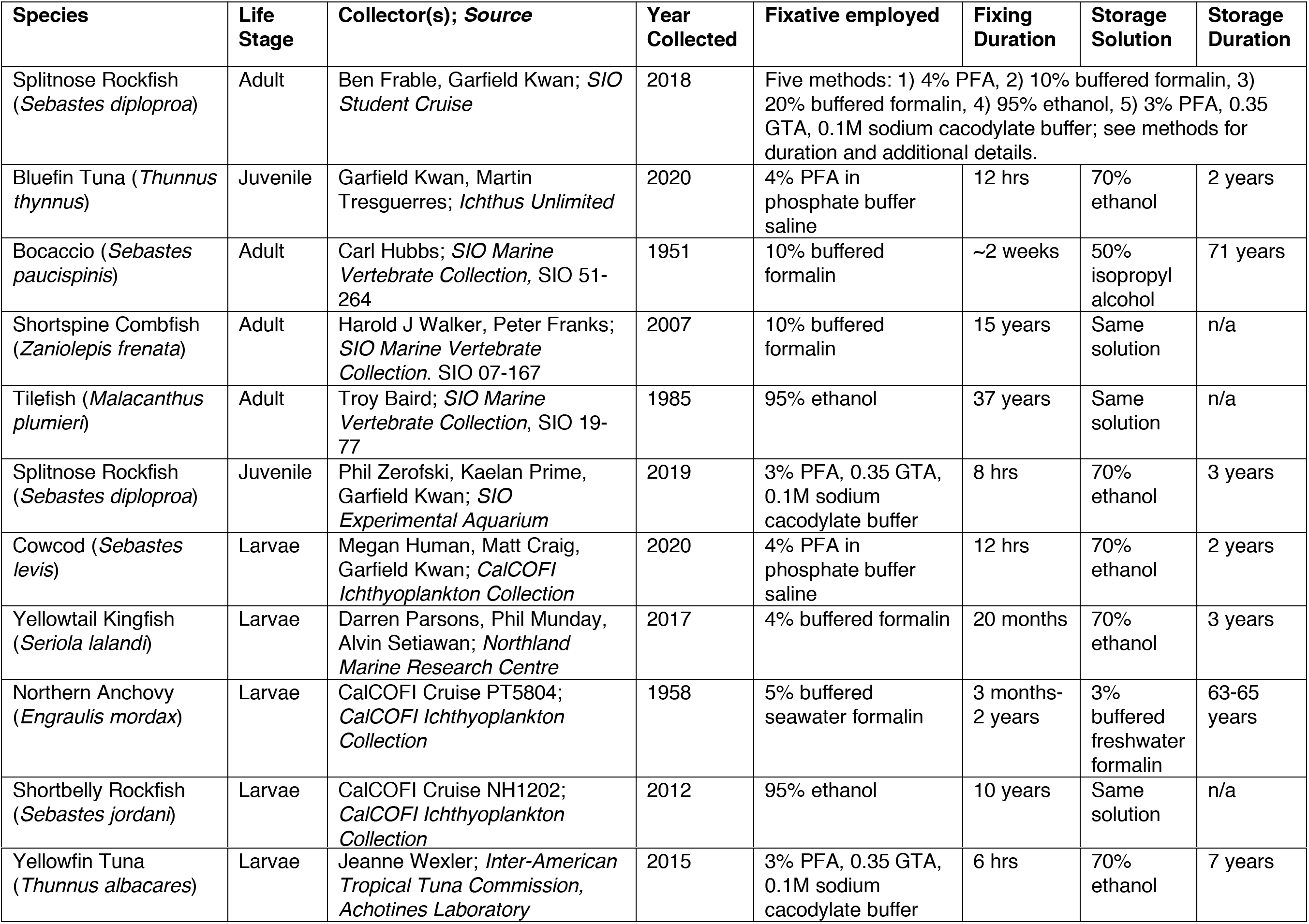
List of fishes tested, and relevant fixation information.

## Methods

### Tissue collection and fixation methods

Splitnose rockfish (*Sebastes diploproa;* N=3) were collected via bottom trawl at 340m depth off Coronado Island, San Diego, California, USA (trawl in: 32 °40.88’N, 117° 23.5’W; trawl out: 32° 43.27’N, 117° 22.09’W) on May 12^th^, 2018. Rockfish gills were quickly extracted and preserved with one of five fixation methods: **1)** 4% PFA in phosphate buffer saline (PBS) at 4°C for 8 hrs, 50% ethanol at 4°C for 12 hrs, then 70% ethanol at 4°C until processing, **2)** 10% formalin buffered with sodium tetraborate at room temperature for 2 weeks, then 50% isopropyl at room temperature until processing, **3)** 20% formalin buffered with sodium tetraborate at room temperature for 2 weeks, then 50% isopropyl at room temperature until processing, **4)** 95% ethanol at 4°C for 8 hrs, then switched to fresh 95% ethanol at room temperature until processing, or **5)** 3% PFA, 0.35% GTA in 0.1M sodium cacodylate buffer at 4°C for 8 hrs, 50% ethanol at 4°C for 12 hrs, then 70% ethanol at 4°C until processing. Processing began ~3 months later on August 15^th^, 2018. For the sake of brevity, we will refer to the preservative “3% PFA, 0.35% GTA in 0.1M sodium cacodylate buffer” simply as *“SEM fixative”*.

### Tissue embedding, sectioning, and epifluorescence imaging

Samples were processed following the protocol detailed in Kwan *et al*. (2020). Gill samples were dehydrated in ethanol (70%, 95%, 100%, 10 min each), incubated in SafeClear (three times; 10 min each), immersed in warm paraffin (65°C; three times; 10 min each), then embedded in paraffin on an ice pack overnight. The next day, gill samples were sectioned using a microtome (12 μm thickness), mounted onto glass slides, dried overnight in an incubator (35°C), and stored at room temperature.

On the day of imaging, paraffin was removed via SafeClear washes (three times; 10 min each), and rehydrated in a series of decreasing ethanol steps (100%, 95%, 70%, 10 min each). Next, samples were subjected to one of the following four quenching methods: **1)** control (PBS), **2)** SBH (1.0 mg/ml) in ice cold PBS (six times; 10 min each), **3)** bath in heated CB (~95°C for 15 min), or **4)** CB then SBH quenching procedure. Samples were then nuclear stained with Hoechst 33342 (5 μg/ml) at room temperature for 1 hr, washed thrice with PBS with 0.1% Tween (PBS-T), immersed in Fluoro-gel with Tris (Electron Microscopy Sciences; Hatfield, Pennsylvania, USA), mounted with a coverslip, and imaged on an epifluorescence microscope (Zeiss AxioObserver Z1; Oberkochen, Germany) equipped with filter cubes Zeiss 49 DAPI (excitation: 352 nm, emission: 455 nm), Zeiss 38 HE GFP (excitation: 493 nm, emission: 517 nm), and Zeiss 43 HE DsRed (excitation: 557 nm, emission: 572 nm) at 20x magnification (objective: Plan-Apochromat 20x/0.8 M27).

Immunostained images were captured using Zeiss Axiovision software at the same exposure duration (DAPI: 50 ms, GFP and DsRed: 1 sec), and their brightness and contrast were not adjusted. Immunostained images were captured using Zeiss Axiovision software at the same exposure duration (DAPI: 50 ms, GFP and DsRed: 1 sec), and their brightness and contrast were not adjusted. DAPI staining was excited at 335-383 nm, and detected at 420-470 nm. Fluorescence in the *green* channel was excited at 450-490 nm, and detected at 500-550 nm. Finally, fluorescence in the *red* channel was excited at 538-562 nm and detected at 570-640 nm.

### Quantifying mean fluorescence intensity

Gill images were separated into green and red channels, converted to greyscale, then imported into FIJI (Schindelin et al., 2012). The fluorescence intensity of both gill filament and background signal were thrice measured (4 x 4μm; 16 μm^2^ per measurement) using the mean fluorescence intensity (MFI) quantification methods detailed in Shihan *et al*. (2021). To ensure sampling consistency, we sampled the gill filament and ensured no background space was present. Autofluorescence was calculated by subtracting gill filament signal from background signal (thrice measured, 4 x 4μm; 16 μm^2^ per measurement). MFI values are presented as a percentage by dividing from 255, the maximum value possible. In total, we examined three gill samples per combination of fixative and quenching method.

### Validating IHC on archival samples

Whole-mount immunostaining using fluorescent secondary antibodies were detailed in Kwan and Tresguerres (2022), respectively. Comparison of quenching techniques across gill sections revealed SBH was the best treatment (see Results), and was selected as the method used to validate archival samples. In anticipation of higher background signal due to longer fixation period, the concentration and number of SBH washes were increased to 1.5 mg/mL and twelve times at 10 min each, respectively. Next, samples were washed with PBS-T at room temperature for 5 min, immersed in blocking buffer (PBS-T, 2% normal goat serum, 0.02% keyhole limpet hemocyanin) at room temperature for 1 hr, then incubated with the primary antibodies (a5: 42 ng/mL in blocking buffer; Developmental Studies Hybridoma Bank, University of Iowa, Iowa City, USA) in blocking buffer at 4°C overnight. The next day, samples were washed in PBS-T (three times; 10 min each), incubated with anti-mouse fluorescent secondary antibodies (Alexa Fluor 545; 1:1,000), anti-rabbit fluorescent secondary antibodies (Alexa Fluor 488; 1:1000) and Hoechst 33342 (5 μg/ml; Invitrogen) at room temperature for 1 hr, washed in PBS-T (three times; 10 min each), then mounted onto depression slides with a cover slip.

AlexaFluor (to visualize NKA) was detected in archival samples using the Zeiss AxioObserver Z1 with a 20x objective (Plan-Apochromat 20x/0.8 M27). Samples were imaged for a total of four times: pre-SBH treatment (epifluorescence, DAPI, GFP, and DsRed: 100 ms), post-SBH treatment (epifluorescence, DAPI, GFP, and DsRed: 100 ms), post-immunostaining epifluorescence microscopy (epifluorescence, DAPI: 3 ms, GFP and DsRed: 40 ms), and post-immunostaining confocal microscopy (confocal, DAPI: 353 nm excitation at 0.8% laser power, 465 nm emission, detection 410-470 nm; GFP: 493 nm excitation at 0.8% laser power, 517 nm emission, detection 510-575 nm; DsRed: 577 nm excitation at 0.8% laser power, 603 nm emission, detection 575-617 nm). Samples without primary antibodies (negative controls) revealed background fluorescence, but no specific ionocyte staining. Adult fishes were imaged on the trailing edge of the gill, whereas larval fishes were imaged on their skin immediately posterior to their respective operculum. Both post-immunostaining epifluorescence and confocal microscope images were Z-stacked imaged at the same location. Image brightness and contrast were not adjusted.

### Statistical analysis

Statistical tests were performed using R (version 4.0.3; R Development Core Team, 2013). Shapiro-Wilks and Levene’s test were used initially to test ANOVA’s assumption of normality and homoscedasticity of variance. First, we ran Two-way ANOVA to evaluate the multiplicative effects of fluorescence channel and fixative on gill sections prior to quenching treatments. We followed this with Tukey’s Honest Significant Difference (HSD) post-hoc to evaluate the significance of pairwise treatments. Second, we again used Two-way ANOVAs and Tukey HSDs tests to elucidate the effects of fixative and quenching. Because our first analysis (see Results) indicated differences between the green and red fluorescence, the two channels were evaluated as separate Two-way ANOVAs.

## Result and Discussion

### Comparison of fixative-induced autofluorescence

Shapiro-Wilks and Levene’s tests indicated that normality and variance assumptions were met for all comparisons. The interaction between fluorescence channel and fixative (F_4,20_=30.99, p=<0.0001) was significant among control, non-quenched samples (Two-way ANOVA; Supplemental Table 1). In addition, green MFI was significantly higher than red in all fixatives except 4% PFA (Tukey HSD; Figure 1). In the green channel, the autofluorescence of tissues fixed in 10% formalin, 20% formalin, and SEM fixative were significantly higher than those preserved with 4% PFA (Figure 1). However, the autofluorescence of tissues fixed in 95% ethanol was not significantly different from those preserved with 4% PFA (Figure 1). In contrast, the type of fixative did not affect autofluorescence in the red channel (Figure 1). Altogether, these results suggest fixative-induced autofluorescence are channel/wavelength specific. Representative images of non-quenched rockfish gill tissues are shown in Figures 2 and 3, respectively.

**Figure 1.**
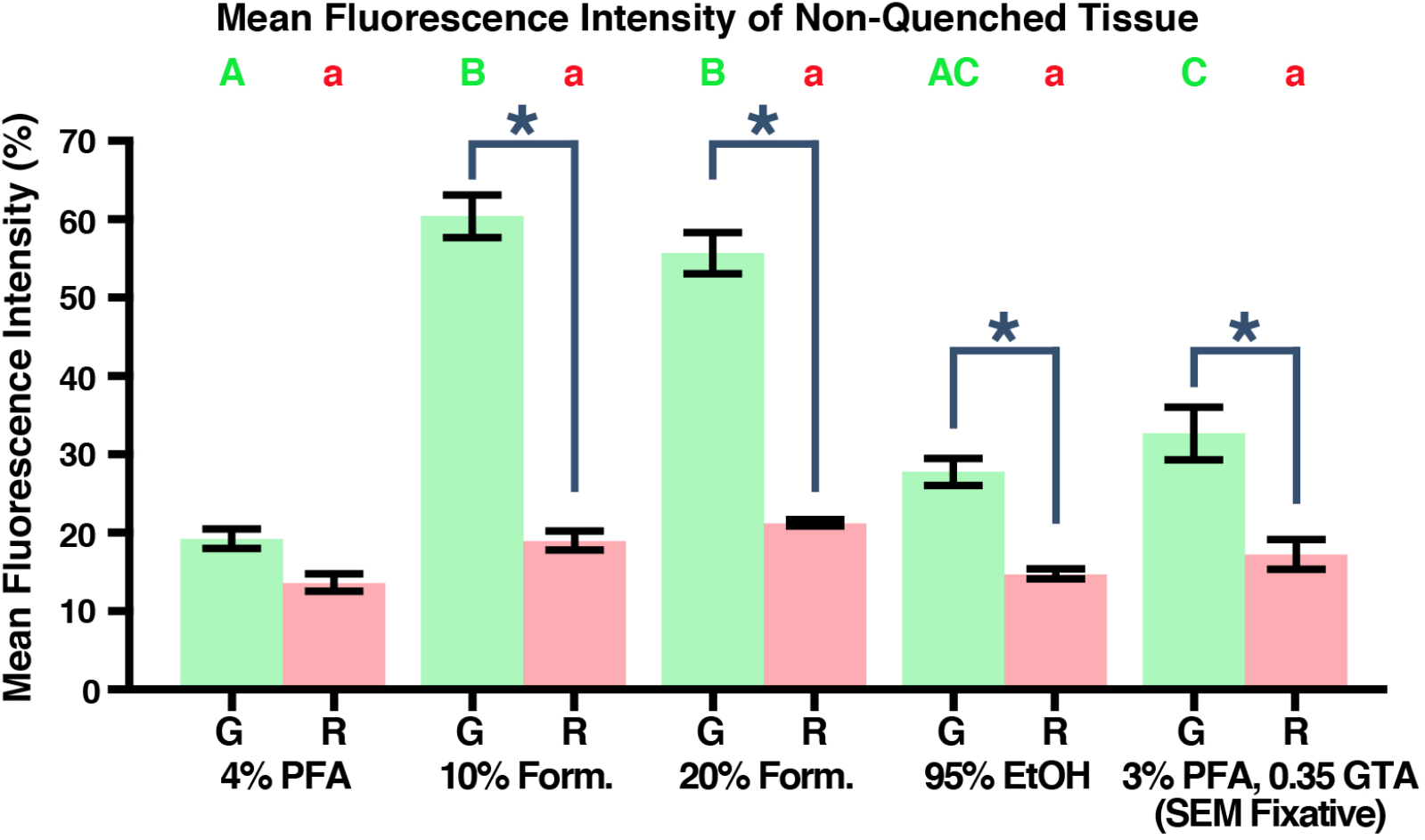
Comparison of fixative-induced autofluorescence prior to quenching treatments in A) green and B) red channels. *Statistical analysis:* 2-way ANOVA with Tukey HSD. Results shown as mean ± SE. *Abbreviations:* G = green, R = red, PFA = paraformaldehyde, Form. = formalin, EtOH = ethanol, GTA = glutaraldehyde. Uppercased green letters denote significance among fixatives within green fluorescence channels, and lowercased red letters denote significance among fixatives within red fluorescence channels. Asterisk denote significance between fluorescence channels within fixatives.

**Figure 2:**
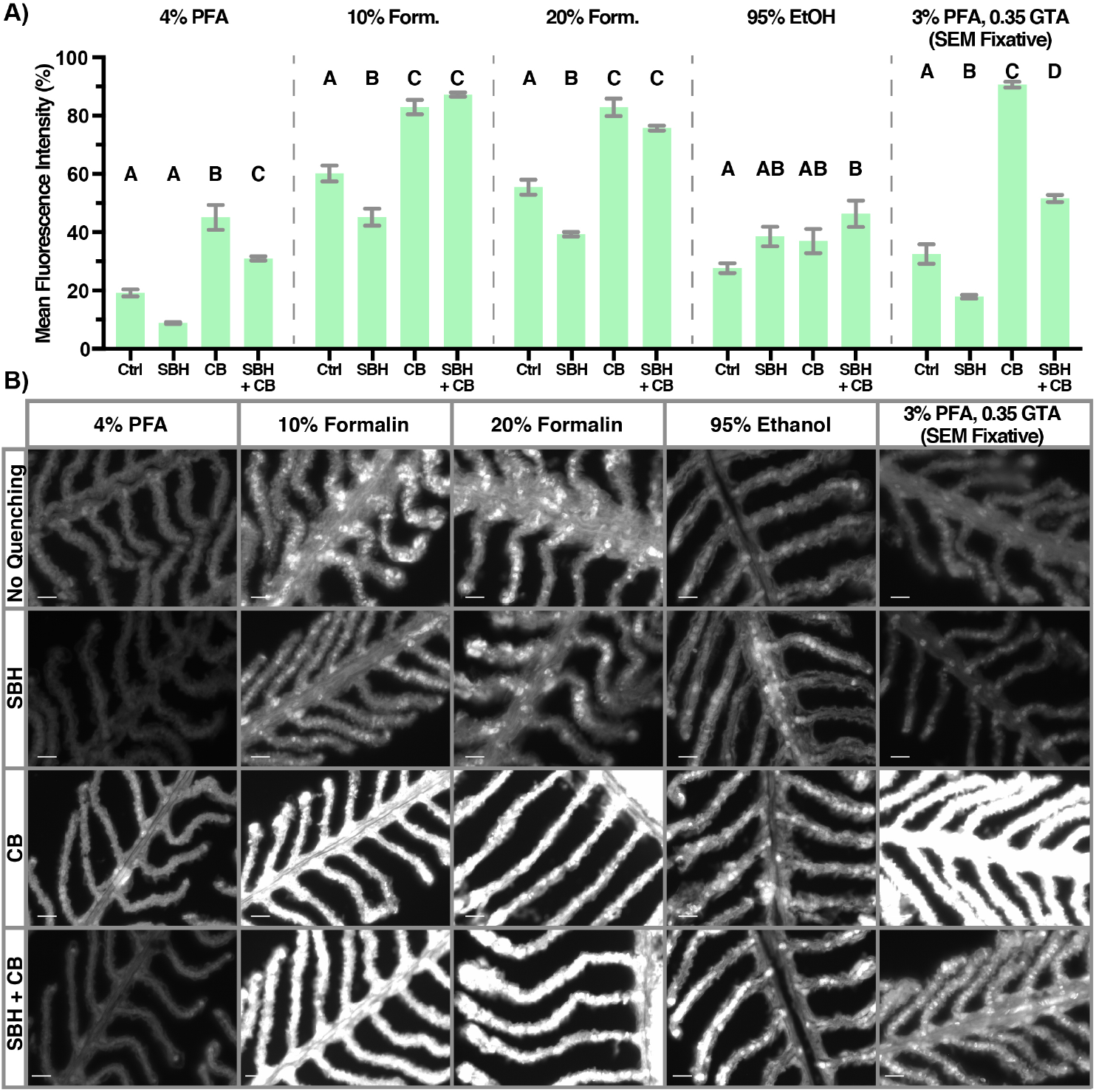
A) Comparison of quenching treatments on fixative-specific autofluorescence in the green channel (excitation: 450-490 nm, detection: 500-550 nm), and B) representative image of gill autofluorescence. All images were captured using the same settings, and none of the images were altered. Alphabet letters denote significance among quenching treatment within fixative. *Statistical analysis:* Two-way ANOVA with Tukey HSD. Results shown as mean ± SE. Scale bar = 20 μm. *Abbreviation:* Ctrl = control (no quenching), PFA = paraformaldehyde, GTA = glutaraldehyde, Form. = formalin, EtOH = ethanol, SBH = sodium borohydride, CB = citrate buffer.

**Figure 3:**
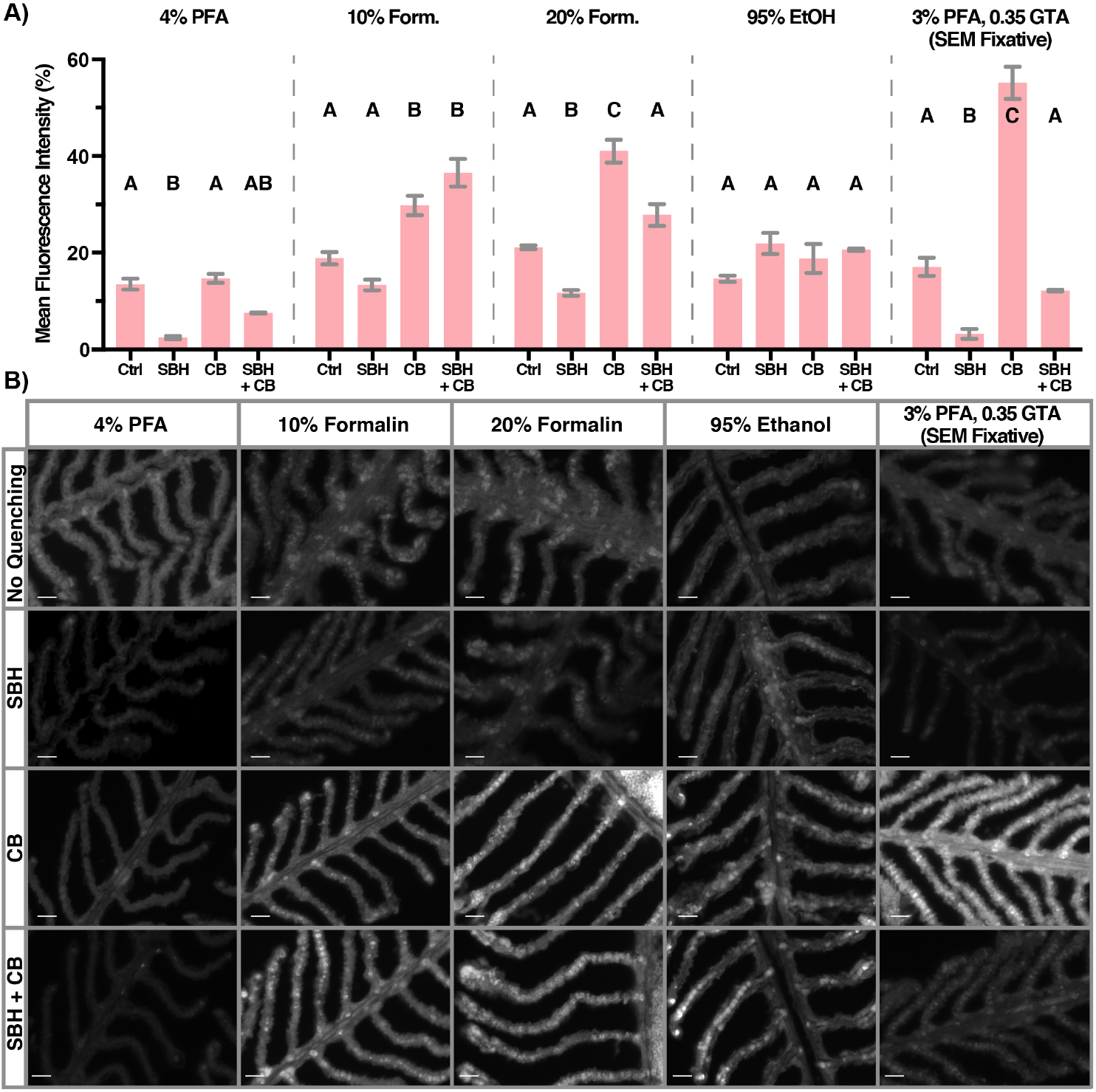
A) Comparison of quenching treatments on fixative-specific autofluorescence in the red channel (excitation: 538-562 nm, detection: 570–640 nm), and B) representative image of gill autofluorescence. All images were captured using the same settings, and none of the images were altered. Alphabet letters denote significance among quenching treatment within fixative. *Statistical analysis:* Two-way ANOVA with Tukey HSD. Results shown as mean ± SE. Scale bar = 20 μm. *Abbreviation:* Ctrl = control (no quenching), PFA = paraformaldehyde, GTA = glutaraldehyde, Form. = formalin, EtOH = ethanol, SBH = sodium borohydride, CB = citrate buffer.

### Impact of quenching on fixative-induced autofluorescence

Due to the disparity in green and red MFI within non-quenched samples, we conducted separate Two-way ANOVAS to analyze the effects of quenching on fixative-induced autofluorescence. For green autoflorescence, the interaction between fixative and quenching methods was highly significant (Two-way ANOVA; F_12,40_=22.83, p=<0.0001; Supplemental Table 2), and varied greatly from 9% (4% PFA and SBH) to 91% (SEM fixative and CB). In general, PFA-, 10% and 20% formalin-, and SEM-fixed samples exhibited significantly reduced green autofluorescence following SBH quenching, and significantly greater green autofluorescence after CB or combined CB and SBH quenching (Figure 2). The only exception was found in 4% PFA-fixed samples: SBH quenching reduced green autofluorescence by ~10% MFI (~50% of control value), but statistical significance was not detected (Figure 2). On the other hand, 95% ethanol-fixed samples were not affected by SBH and CB, but experienced greater green autofluorescence following combined SBH + CB incubation (Figure 2).

For red autofluorescence, the interaction between fixative and quenching methods was also highly significant (Two-way ANOVA; F_12,40_=36.61, p=<0.0001; Supplemental Table 3) and varied from 2% (4% PFA and SBH) to 55% (SEM fixative and CB). The direction in which red autofluorescence responds to the various quenching treatments was mostly consistent with those observed for green autofluorescence. Tukey’s HSD indicated SBH quenching significantly reduced red autofluorescence in 4% PFA-, 20% formalin-, and SEM-fixed samples, but did not significantly affect 10% formalin-and 95% ethanol-fixed samples (Figure 3). Next, CB quenching significantly increased red autofluorescence in 10% and 20% formalin- and SEM-fixed samples, but did not significantly affect samples preserved with 4% PFA and 95% ethanol (Figure 3). Finally, combined SBH and CB resulted in significantly higher red autofluorescence in 10% formalin fixed samples, and did not significantly change samples preserved in 4% PFA, 20% formalin, 95% ethanol, or SEM fixative (Figure 3).

In summary, SBH incubation generally reduced a considerable amount of red and green autofluorescence in PFA-, formalin- and GTA-fixed samples. Because SBH is a known aldehydic reducing agent (Abdel-Akher et al., 1952), its capacity to reduce cross-linkage caused by aldehyde fixatives (e.g. PFA, formalin, GTA) (Thavarajah et al., 2012) should not be surprising. Moreover, this may explain why samples preserved with 95% ethanol failed to improve following SBH incubation since their observed autofluorescence were not derived from the aldehyde. Finally, quenching with solely CB or in combination with SBH was rarely effective at reducing autofluorescence, thereby suggesting high temperature and alkaline hydrolysis are not appropriate for reducing background signal in aldehyde- and ethanol-preserved archival fish samples.

### Validating fluorescent IHC on whole-mount archival samples

Because SBH showed the most potential in reducing autofluorescence, we focused on validating this quenching treatment on whole-mount archival samples. We selected archived fish samples that vary in species, fixative, and fixation duration to determine if IHC is viable for real-world application within natural history collections (Table 1). In addition, we chose NKA as our protein-of-interest due to the many past IHC studies that successfully documented this protein in larval fish skin (reviewed in Varsamos et al., 2005) and adult fish gills (reviewed in Hwang and Lee, 2007; Hwang et al., 2011). Consistent with our earlier findings, SBH incubation dramatically reduced autofluorescence in whole-mount samples fixed with PFA, formalin, and GTA – but not in samples fixed with 95% ethanol (Supplemental Figures 1, 2). Moreover, NKA was successfully immunolocalized within skin ionocytes of larval fishes and gill ionocytes of juvenile and adult fishes as observed in past IHC studies.

Next, we assessed whether we could visualize NKA within the skin and gill of various archival samples using epifluorescence microscopy (Figure 4). As expected, NKA-rich ionocytes were most discernable in samples with low background signals such as the PFA-fixed cowcod and bluefin tuna. We also successfully increased the signal-to-noise ratio within formalin-fixed yellowtail kingfish (formalin fixed for 20 months) to levels comparable to the PFA-fixed samples. However, this level of optimization was not consistently observed across formalin-fixed samples. For instance, NKA signal was not detectable in the anchovy (formalin fixed for 71 years) despite relatively low autofluorescence, whereas NKA signal within bocaccio (formalin fixed for 2 weeks) and shortspine combfish (formalin fixed for 15 years) were somewhat visible despite relatively high autofluorescence. This suggests the duration of formalin fixation affects the signal-to-noise ratio. Samples fixed with 95% ethanol had intensely bright autofluorescence, drowning out any NKA signal that may have been present in the shortbelly rockfish and tilefish. Conversely, despite relatively low autofluorescence, NKA signals were not observed in GTA-fixed yellowfin tuna and splitnose rockfish.

**Figure 4:**
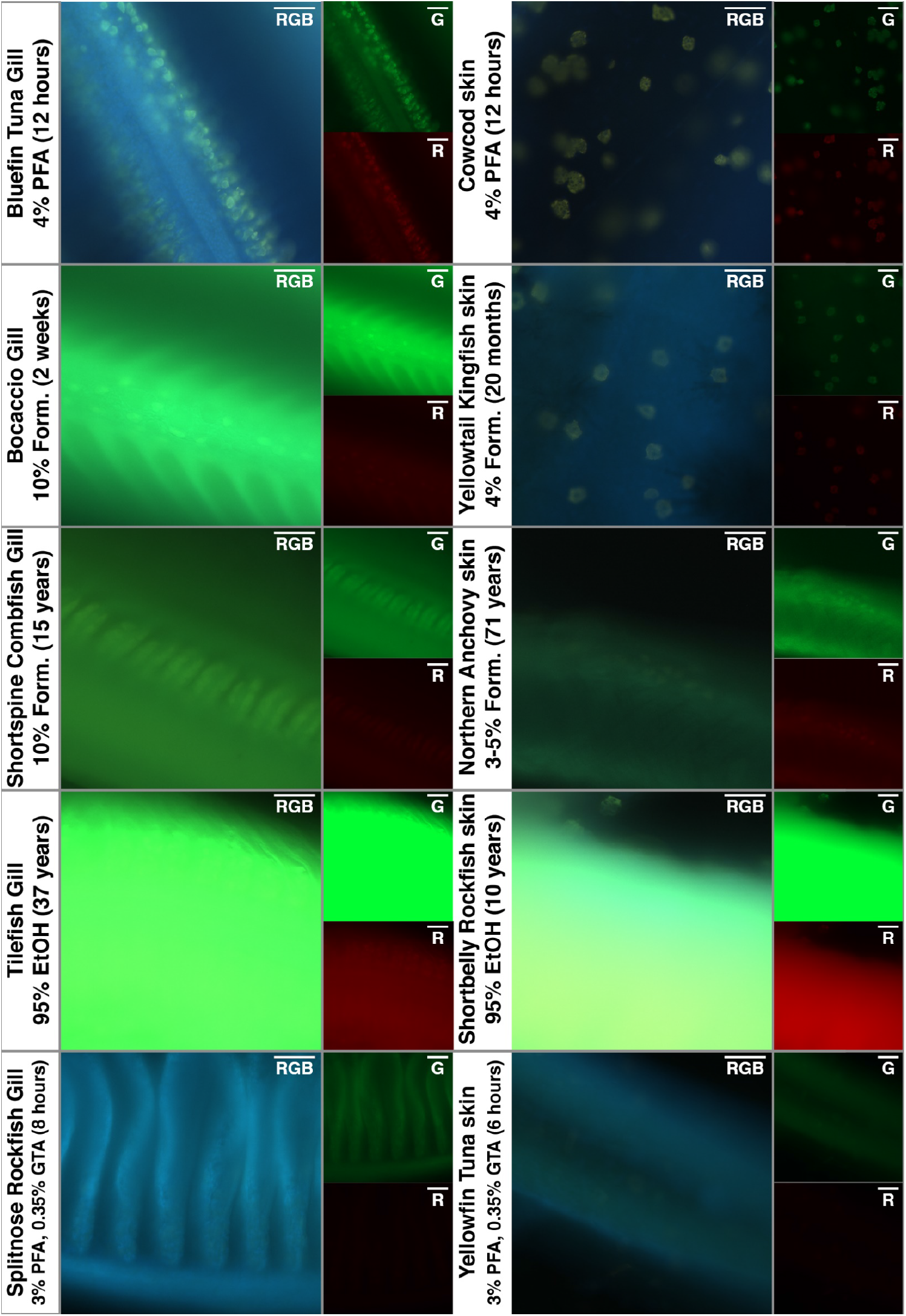
NKA (green, red) within fish samples preserved with various fixatives and fixation duration were whole-mount imaged with epifluorescence microscopy. Images show a single image as high background signal prevents maximum intensity projection. Brightness and contrast were not adjusted.

Unlike epifluorescence microscopy, confocal microscopy can filter the signal from the background and out-of-focus noise. As expected, confocal microscopy greatly enhanced the signal-to-noise ratio, allowing us to identify NKA-rich ionocytes despite their high autofluorescence and/or dim signal regardless of fixative and fixation duration (Figure 5). In PFA-fixed cowcod and bluefin tuna samples, NKA signal were distinct and greatly enhanced. This sharp NKA signal was also observed in samples that were formalin-fixed for a relatively short period (bocaccio, 2 weeks; yellowtail kingfish, 20 months), and rivals the quality of PFA-fixed samples. Confocal microscopy also greatly enhanced NKA signal in samples that were formalin-fixed for longer durations (shortspine combfish, 15 years; anchovy, 63-65 years), though their background signals remained apparent. In addition, confocal microscopy was able to distinguish the NKA signal within ethanol-fixed anchovy and tilefish (albeit with high background signal) and in GTA-fixed yellowfin tuna and splitnose rockfish. These results demonstrate the combination of SBH quenching and confocal microscopy could be used to successfully immunotarget archival samples with fluorescent probes, and produce images of higher quality compared to those captured with epifluorescence microscopy.

**Figure 5:**
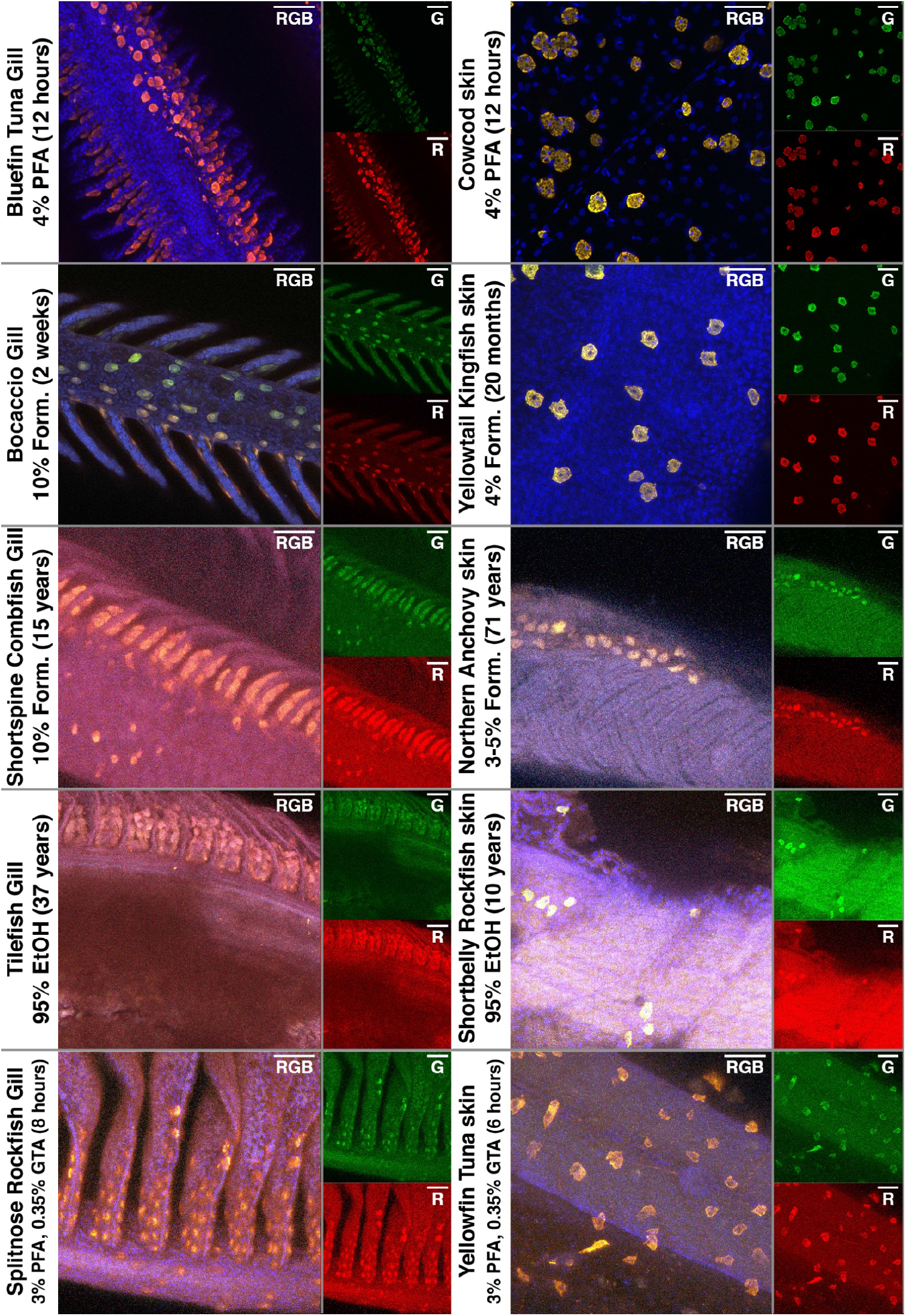
NKA (green, red) within fish samples preserved with various fixatives and fixation duration were whole-mount imaged with confocal microscopy. Images are shown as maximum intensity projection. Brightness and contrast were not adjusted.

### Implications for natural history collections

In this study, we demonstrated the feasibility of fluorescent immunostaining of archival samples, including those that have been immersed within formalin- or ethanol for multiple decades. We found repeated SBH washes greatly reduced autofluorescence in tissues preserved with aldehyde-based fixatives, and greatly enhanced downstream immunostaining and imaging. While epifluorescence imaging could not adequately detect our protein-of-interest within formalin-, ethanol-, and GTA-fixed samples, confocal microscopy was able to resolve the NKA signal from the background in both red and green channels, regardless of fixative and fixation duration. Altogether, this study suggests archival samples housed within natural history collections can be immunostained with relatively high success.

The duration of formalin fixation ranges from weeks to years depending on specimen size, collection protocols, and other circumstances (e.g. lack of resources at field station, time collected on sampling cruises). Here, we demonstrated that formalin-fixed samples that were transferred into isopropyl after ~2-weeks (bocaccio) and into ethanol after 2-years’ time (yellowtail kingfish) can be sufficiently quenched to produce IHC at a quality that rival samples preserved with PFA (bluefin tuna, cowcod). In particular, the bocaccio sample has been stored within isopropyl for ~71 years, and the quality of the resulting IHC images are particularly encouraging as many archival samples are eventually transferred into alcohol for long-term storage once the specimen is thought to be properly preserved.

Natural history collections sometimes opt to keep specimens within formalin to avoid the shrinkage effect associated with ethanol preservation. We observed extended formalin fixation resulted in higher autofluorescence and obvious differences in IHC quality when compared to the formalin-fixed specimens that were eventually transferred into alcohol (e.g. bocaccio and yellowtail kingfish). Even so, the signal-to-noise ratio was sufficiently high enough to identify the NKA signal within samples fixed in formalin for 15 (shortspine combfish) and 65 years (Northern anchovy). These results demonstrate specimens that have been completely formalin-immersed can still be viable for IHC. To the best of our knowledge, this study is the first to have successfully immunostained a specimen that was formalin preserved for 65 years.

Although ethanol- and GTA-fixed samples are not typically used for IHC, we included these two treatments to demonstrate their capacity to serve as a last resort. Although SBH had little impact on the autofluorescence of ethanol-fixed samples, confocal microscopy was still capable of immunostaining our protein-of-interest. In contrast, although SBH reduced autofluorescence in GTA-fixed samples, these still exhibited high background autofluorescence, and the resulting quality was similar to those fixed with ethanol and formalin-only specimens.

Due to the comparative nature of this study, we opted to restrict imaging parameters including exposure duration, laser intensity, and channel-specific brightness and contrast levels. Needless to say, sample-specific adjustments would have enhanced the image quality of specimens fixed with non-optimal preservation methods. Researchers interested in fluorescently immunostaining archival samples should also explore fine-tuning microscope software to adjusting the wavelengths of interest, and to utilize fluorophores that emit at wavelengths with low autofluorescence. In addition, modern confocal microscope software has the capacity to subtract specific wavelengths, which can also be used to reduce autofluorescence. Suffice to say, there are many options to improve the quality of fluorescent IHC in problematic samples. On another note, many of these techniques, such as SBH’s ability to quench autofluorescence, may be applicable for improving other protocols such as fluorescent *in situ* hybridization (FISH), especially in ethanol-fixed, DNA/RNA stable specimens (Oliveira et al., 2010; Benerini Gatta et al., 2012).

By optimizing IHC on archival samples, we provide a valuable tool to examine biological responses across archives collected from decades to centuries ago. For instance, we focused on immunostaining NKA-rich ionocytes, which are ion-transporting cells responsible for maintaining osmotic and acid-base homeostasis (reviewed in Evans et al., 2005; Marshall and Grosell, 2006). As climate change progresses and aquatic habitats are modified due to anthropogenic activities, the immunostaining of gill ionocytes can be used to establish baselines as comparison, and to better predict future physiological responses of teleost fishes. For instance, past studies showed that the type of protein transporters synthesized within NKA-rich ionocytes are dependent on the salinity of the environment (reviewed in Hiroi and McCormick, 2012), and additional NKA-rich ionocytes may form in response to hyper- (Varsamos et al., 2002b) and hypo-osmotic stress (Uchida and Kaneko, 1996; Sasai et al., 1998; Hirai et al., 1999; Zydlewski and McCormick, 2001). Today, freshwater rivers are increasingly diverted for use in farmlands, dams, and cities (e.g. San Francisco Estuary), estuarine environments are becoming more saline due to saltwater intrusion (Cloern et al., 2011; Hutton et al., 2016). This increase in salinity may result in changes in protein expression and ionocyte abundance, and can be elucidated using the described IHC and quenching techniques to compare archived estuarine fishes collected from decades ago to their contemporary counterparts. Similarly, IHC on archived fish specimens can be applied to other aquatic habitats to assess the biological responses of climate change across decadal and centurial timescales.

Finally, while we focused on teleost tissue, these IHC techniques can be applied to other organisms (e.g. annelid, arthropod, mollusk) archived within natural history collections, ultimately allowing for the study of tissues from historical, endangered, and extinct species. However, and as shown here, some optimization is to be expected. Besides SBH, other quenching strategies previously reported to reduce autofluorescence and may be worth exploring include photobleaching with UV irradiation, ammonia + ethanol washes, and staining with dyes such as Sudan Black B (Ramos-Vara, 2005; Oliveira et al., 2010). Altogether, this study showcases the potential for using IHC facilitated with SBH to analyze specimens collected from decades to centuries ago – thereby providing researchers with another tool to explore biological responses over time, and by extension, elevate the inherent value of natural history collections.

## Acknowledgement

GTK was funded by the NSF GRFP and NSF PRFB (Award #1907334). This project was also supported by SIO discretionary fund awarded to MT. We thank Dr. Jennifer Taylor and the R/V Robert Gordon Sproul crew members for the opportunity to collect rockfish. We are indebted to the *SIO Marine Vertebrate Collection* and the *CalCOFI Ichthyoplankton Collection* for providing valuable archival specimens, and their respective curators and collection managers for maintaining the natural history collections. We are also grateful to many researchers who collected and contributed specimens for this study, including Dr. Phil Hastings, Harold J. Walker, William Watson, Sherri Charter, Dr. Carl Hubbs, Dr. Peter Franks, Dr. Alejandro Buentello and the Ichthus Unlimited team, Dr. Troy Baird, Kaelan Prime, Dr. Darren Parsons, Dr. Phil Munday, Dr. Alvin Setiawan, the Northland Marine Research Centre team, Megan Human, Dr. Matt Craig, Jeanne Wexler, the technical staff at Achotines Laboratory in Panama, Dr. Daniel Margulies, Vernon Scholey, the Inter-American Tropical Tuna Commission, the CalCOFI crew and scientists abroad the R/V Paolina T. on cruise 5804 and R/V New Horizon cruise 1202.

## References

Abdel-Akher M, Hamilton TK, Montgomery R, Smith F (1952) A new procedure for the determination of the fine structure of polysaccharides. J Am Chem Soc 74:4970–4971. doi: 10.1021/ja01139a526

Baschong W, Suetterlin R, Laeng RH (2001) Control of autofluorescence of archival formaldehyde-fixed, paraffin-embedded tissue in confocal laser scanning microscopy (CLSM). J Histochem Cytochem 49:1565–1572. doi: 10.1177/002215540104901210

Benerini Gatta L, Cadei M, Balzarini P, et al (2012) Application of alternative fixatives to formalin in diagnostic pathology. Eur J Histochem 56:63–70. doi: 10.4081/ejh.2012.e12

Card DC, Shapiro B, Giribet G, et al (2021) Museum Genomics. Annu Rev Genet 55:633–659. doi: 10.1146/annurev-genet-071719-020506

Christensen AK, Hiroi J, Schultz ET, McCormick SD (2012) Branchial ionocyte organization and ion-transport protein expression in juvenile alewives acclimated to freshwater or seawater. J Exp Biol 215:642–652. doi: 10.1242/jeb.063057

Clancy B, Cauller LJ (1998) Reduction of background autofluorescence in br.pdf. 83:97–102.

Cloern JE, Knowles N, Brown LR, et al (2011) Projected evolution of California’s San Francisco bay-delta-river system in a century of climate change. PLoS One. doi: 10.1371/journal.pone.0024465

Diez-Fraile A, Van N, J. C, DHerde K (2012) Optimizing Multiple Immunostaining of Neural Tissue. In: Applications of Immunocytochemistry. pp 345–442

Edwards A, Melanie S, Thomas F (2002) Short-and Long-Term Effects of Fixation and Preservation on Stable Isotope Values (δ13C, δ15N, δ34S) of Fluid-Preserved Museum Specimens. Copeia 4:1106–1112. doi: 10.1643/0045-8511(2002)002[1106:SALTEO]2.0.CO;2

Esbaugh AJ, Cutler B (2016) Intestinal Na+, K+, 2Cl-cotransporter 2 plays a crucial role in hyperosmotic transitions of a euryhaline teleost. Physiol Rep 4:1–12. doi: 10.14814/phy2.13028

Evans DH, Piermarini PM, Choe KP (2005) The Multifunctional Fish Gill: Dominant Site of Gas Exchange, Osmoregulation, Acid-Base Regulation, and Excretion of Nitrogenous Waste. Physiol Rev 85:97–177. doi: 10.1152/physrev.00050.2003

Frable BW (2022) Personal Communication: About the Marine Vertebrate Collection. https://scripps.ucsd.edu/marine-vertebrate-collection/about-marine-vertebrate-collection. Accessed 9 Jun 2022

Frommel AY, Kwan GT, Prime KJ, et al (2021) Changes in gill and air-breathing organ characteristics during the transition from water-to air-breathing in juvenile Arapaima gigas. J Exp Zool Part A Ecol Integr Physiol 335:jez.2456. doi: 10.1002/jez.2456

Glover CN, Bucking C, Wood CM (2013) The skin of fish as a transport epithelium: A review. J Comp Physiol B Biochem Syst Environ Physiol 183:877–891. doi: 10.1007/s00360-013-0761-4

Hetherington ED, Kurle CM, Ohman MD, Popp BN (2019) Effects of chemical preservation on bulk and amino acid isotope ratios of zooplankton, fish, and squid tissues. Rapid Commun Mass Spectrom 33:935–945. doi: 10.1002/rcm.8408

Hirai N, Tagawa M, Kaneko T, et al (1999) Distributional changes in branchial chloride cells during freshwater adaptation in Japanese Sea Bass Lateolabrax japonicus. Zoolog Sci 16:43–49. doi: 10.2108/zsj.16.43

Hiroi J, McCormick SD (2012) New insights into gill ionocyte and ion transporter function in euryhaline and diadromous fish. Respir Physiol Neurobiol 184:257–268. doi: 10.1016/j.resp.2012.07.019

Hutton PH, Rath JS, Chen L, et al (2016) Nine Decades of Salinity Observations in the San Francisco Bay and Delta: Modeling and Trend Evaluations. J Water Resour Plan Manag 142:04015069. doi: 10.1061/(asce)wr.1943-5452.0000617

Hwang P-P, Lee T-H, Lin L-Y (2011) Ion regulation in fish gills: recent progress in the cellular and molecular mechanisms. AJP Regul Integr Comp Physiol 301:R28–R47. doi: 10.1152/ajpregu.00047.2011

Hwang PP, Lee TH (2007) New insights into fish ion regulation and mitochondrion-rich cells. Comp Biochem Physiol - A Mol Integr Physiol 148:479–497. doi: 10.1016/j.cbpa.2007.06.416

Kelly B, Dempson JB, Power M (2006) The effects of preservation on fish tissue stable isotope signatures. J Fish Biol 69:1595–1611. doi: 10.1111/j.1095-8649.2006.01226.x

Kiernan JA (2000) Formaldehyde, Formalin, Paraformaldehyde And Glutaraldehyde: What They Are And What They Do. Micros Today 8:8–13. doi: 10.1017/s1551929500057060

Kwan GT, Wexler JB, Wegner NC, Tresguerres M (2019) Ontogenetic changes in cutaneous and branchial ionocytes and morphology in yellowfin tuna (Thunnus albacares) larvae. J Comp Physiol B Biochem Syst Environ Physiol 189:81–95. doi: 10.1007/s00360-018-1187-9

Kwan GT, Smith TR, Tresguerres M (2020) Immunological characterization of two types of ionocytes in the inner ear epithelium of Pacific Chub Mackerel (Scomber japonicus). J Comp Physiol B 190:419–431. doi: 10.1007/s00360-020-01276-3

Kwan GT, Tresguerres M (2022) Elucidating the acid-base mechanisms underlying otolith overgrowth in fish exposed to ocean acidification. Sci Total Environ 823:153690. doi: 10.1016/j.scitotenv.2022.153690

Luquin E, Pérez-Lorenzo E, Aymerich MS, Mengual E (2010) Two-color fluorescence labeling in acrolein-fixed brain tissue. J Histochem Cytochem 58:359–368. doi: 10.1369/jhc.2009.954495

Marshall WS, Grosell M (2006) Ion Transport, Osmoregulation, and Acid-Base Balance. In: Evans DH, Claiborne JB (eds) The Physiology of Fishes, 3rd edn. CRC Press, Boca Raton, pp 177–230

Matsuda Y, Fujii T, Suzuki T, et al (2011) Comparison of fixation methods for preservation of morphology, RNAs, and proteins from paraffin-embedded human cancer cell-implanted mouse models. J Histochem Cytochem 59:68–75. doi: 10.1369/jhc.2010.957217

Montgomery DW, Kwan GT, Davison WG, et al (2022) Rapid blood acid–base regulation by European sea bass (Dicentrarchus labrax) in response to sudden exposure to high environmental CO2. J Exp Biol. doi: 10.1242/jeb.242735

Moser HG, Charter RL, Smith PE, et al (2002) Distributional atlas of fish larvae and eggs from Manta (surface) samples collected on CalCOFI surveys from 1977 to 2000.

Oliveira VC, Carrara RC V, Simoes DLC, et al (2010) Sudan Black B treatment reduces autofluorescence and improves resolution of in situhybridization specific fluorescent signals of brain sections. Histol Histopathol 25:1017–1024. doi: 10.14670/HH-25.1017

Pikkarainen M, Martikainen P, Alafuzoff I (2010) The effect of prolonged fixation time on immunohistochemical staining of common Neurodegenerative disease markers. J Neuropathol Exp Neurol 69:40–52. doi: 10.1097/NEN.0b013e3181c6c13d

R Development Core Team (2013) R: A language and environment for statistical computing. R foundation for statistical computing.

Ramos-Vara JA (2005) Technical aspects of immunohistochemistry. Vet Pathol 42:405–426. doi: 10.1354/vp.42-4-405

Raxworthy CJ, Smith BT (2021) Mining museums for historical DNA: advances and challenges in museomics. Trends Ecol Evol 36:1049–1060. doi: 10.1016/j.tree.2021.07.009

Sarakinos HC, Johnson ML, Vander Zanden MJ (2002) A synthesis of tissue-preservation effects on carbon and nitrogen stable isotope signatures. Can J Zool 80:381–387. doi: 10.1139/z02-007

Sasai S, Kaneko T, Hasegawa S, Tsukamoto K (1998) Morphological alteration in two types of gill chloride cells in Japanese eels (Anguilla japonica) during catadromous migration. Can J Zool 76:1480–1487. doi: 10.1139/z98-072

Schindelin J, Arganda-Carreras I, Frise E, et al (2012) Fiji: an open-source platform for biological-image analysis. Nat Methods 9:676–682. doi: 10.1038/nmeth.2019

Shaffer HB, Fisher RN, Davidson C (1998) The role of natural history collections in documenting species declines. Trends Ecol Evol 13:27–30. doi: 10.1016/S0169-5347(97)01177-4

Shi S, Cote RJ, Taylor CR (1997) Antigen Retrieval Immunohistochemistry: Past, Present, and Future. J Histochem Cytochem 45:327–343.

Shi SR, Chaiwun B, Young L, et al (1993) Antigen retrieval technique utilizing citrate buffer or urea solution for immunohistochemical demonstration of androgen receptor in formalin-fixed paraffin sections. J Histochem Cytochem 41:1599–1604. doi: 10.1177/41.11.7691930

Shi SR, Shi Y, Taylor CR (2011) Antigen retrieval immunohistochemistry: Review and future prospects in research and diagnosis over two decades. J Histochem Cytochem 59:13–32. doi: 10.1369/jhc.2010.957191

Shihan MH, Novo SG, Le Marchand SJ, et al (2021) A simple method for quantitating confocal fluorescent images. Biochem Biophys Reports 25:100916. doi: 10.1016/j.bbrep.2021.100916

Simmons JE (2014) Fluid Preservation. Rowman & Littlefield, Plymouth, United Kingdom

Singer RA, Love KJ, Page LM (2018) A survey of digitized data from u.S. Fish collections in the idigbio data aggregator. PLoS One 13:1–20. doi: 10.1371/journal.pone.0207636

Stollar D, Grossman L (1962) The reaction of formaldehyde with denatured DNA: Spectrophotometric, immunologic, and enzymic studies. J Mol Biol 4:31–38. doi: 10.1016/S0022-2836(62)80114-4

Stradleigh TW, Ishida AT (2015) Fixation strategies for retinal immunohistochemistry. Prog Retin Eye Res 48:181–202. doi: 10.1016/j.preteyeres.2015.04.001

Swalethorp R, Nielsen TG, Thompson AR, et al (2016) Early life of an inshore population of West Greenlandic cod Gadus morhua: Spatial and temporal aspects of growth and survival. Mar Ecol Prog Ser 555:185–202. doi: 10.3354/meps11816

Thavarajah R, Mudimbaimannar VK, Elizabeth J, et al (2012) Chemical and physical basics of routine formaldehyde fixation. J Oral Maxillofac Pathol 16:400–405. doi: 10.4103/0973-029X.102496

Thompson AR, Chen DC, Guo LW, et al (2017) Larval abundances of rockfishes that were historically targeted by fishing increased over 16 years in association with a large marine protected area. R Soc Open Sci. doi: 10.1098/rsos.170639

Uchida K, Kaneko T (1996) Enhanced chloride cell turnover in the gills of Chum Salmon fry in seawater. Zoolog Sci 13:655–660. doi: 10.2108/zsj.13.655

Varsamos S, Diaz J, Charmantier G, et al (2002a) Location and morphology of chloride cells during the post-embryonic development of the European sea bass, Dicentrarchus labrax. Anat Embryol (Berl) 205:203–213. doi: 10.1007/s00429-002-0231-3

Varsamos S, Diaz JP, Charmantier GUY, et al (2002b) Branchial chloride cells in sea bass (Dicentrarchus labrax) adapted to fresh water, seawater, and doubly concentrated seawater. J Exp Zool 293:12–26. doi: 10.1002/jez.10099

Varsamos S, Nebel C, Charmantier G (2005) Ontogeny of osmoregulation in postembryonic fish: A review. Comp Biochem Physiol - A Mol Integr Physiol 141:401–429. doi: 10.1016/j.cbpb.2005.01.013

Wilson JM, Randall DJ, Donowitz M, et al (2000) Immunolocalization of ion-transport proteins to branchial epithelium mitochondria-rich cells in the mudskipper (Periophthalmodon schlosseri). J Exp Biol 203:2297–2310.

Yang SH, Kang CK, Kung HN, Lee TH (2014) The lamellae-free-type pseudobranch of the euryhaline milkfish (Chanos chanos) is a Na+, K+-ATPase-abundant organ involved in hypoosmoregulation. Comp Biochem Physiol - A Mol Integr Physiol 170:15–25. doi: 10.1016/j.cbpa.2013.12.018

Zydlewski J, McCormick SD (2001) Developmental and environmental regulation of chloride cells in young American shad, Alosa sapidissima. J Exp Zool 290:73–87.

