## Supplemental Information for "Optimizing immunostaining of archival fish samples to enhance museum collection potential"

### Supplemental Material

**Supplemental Table 1:** Relevant Tukey's HSD statistical details on green autofluorescence ~ fixative*quenching. $\alpha$=0.05.

| Pairwise Comparison | Mean Difference | Lower Hinge | Upper Hinge | Adjusted P Value | Significant? |
|---|---|---|---|---|---|
| 20%f:Green-10%f:Green | -4.6849673 | -14.29998 | 4.9300458 | 0.769763 | No |
| 4%pfa:Green-10%f:Green | -40.979521 | -50.594534 | -31.364508 | 0 | Yes |
| 95%EtOH:Green-10%f:Green | -32.468453 | -42.083466 | -22.85344 | 0 | Yes |
| EMS:Green-10%f:Green | -27.59573 | -37.210743 | -17.980717 | 0.0000001 | Yes |
| EMS:Green-95%EtOH:Green | 4.8727233 | -4.74229 | 14.4877365 | 0.731247 | No |
| 95%EtOH:Green-4%pfa:Green | 8.5110675 | -1.103946 | 18.1260807 | 0.1107787 | No |
| EMS:Green-4%pfa:Green | 13.3837909 | 3.768778 | 22.998804 | 0.0025509 | Yes |
| 4%pfa:Green-20%f:Green | -36.294553 | -45.909567 | -26.67954 | 0 | Yes |
| 95%EtOH:Green-20%f:Green | -27.783486 | -37.398499 | -18.168473 | 0.0000001 | Yes |
| EMS:Green-20%f:Green | -22.910763 | -32.525776 | -13.295749 | 0.0000019 | Yes |
| 20%f:Red-10%f:Red | 2.262963 | -7.35205 | 11.8779761 | 0.9969485 | No |
| 4%pfa:Red-10%f:Red | -5.3539869 | -14.969 | 4.2610262 | 0.6256692 | No |
| 95%EtOH:Red-10%f:Red | -4.232244 | -13.847257 | 5.3827691 | 0.8522834 | No |
| EMS:Red-10%f:Red | -1.7864488 | -11.401462 | 7.8285643 | 0.9995053 | No |
| 4%pfa:Red-20%f:Red | -7.6169499 | -17.231963 | 1.9980633 | 0.1997706 | No |
| 95%EtOH:Red-20%f:Red | -6.495207 | -16.11022 | 3.1198062 | 0.3792554 | No |
| EMS:Red-20%f:Red | -4.0494118 | -13.664425 | 5.5656014 | 0.8804908 | No |
| 95%EtOH:Red-4%pfa:Red | 1.1217429 | -8.49327 | 10.7367561 | 0.9999896 | No |
| EMS:Red-4%pfa:Red | 3.5675381 | -6.047475 | 13.1825513 | 0.9387269 | No |
| EMS:Red-95%EtOH:Red | 2.4457952 | -7.169218 | 12.0608084 | 0.9946261 | No |
| EMS:Red-EMS:Green | -15.463486 | -25.078499 | -5.8484727 | 0.0004805 | Yes |
| 95%EtOH:Red-95%EtOH:Green | -13.036558 | -22.651571 | -3.4215446 | 0.0033762 | Yes |
| 20%f:Red-20%f:Green | -34.324837 | -43.93985 | -24.709824 | 0 | Yes |
| 10%f:Red-10%f:Green | -41.272767 | -50.88778 | -31.657754 | 0 | Yes |
| 4%pfa:Red-4%pfa:Green | -5.6472331 | -15.262246 | 3.96778 | 0.5595127 | No |

**Supplemental Table 2:** Relevant Tukey's HSD statistical details on green autofluorescence ~ fixative*quenching. $\alpha$=0.05.

| Pairwise Comparison | Mean Difference | Lower Hinge | Upper Hinge | Adjusted P Value | Significant? |
|---|---|---|---|---|---|
| 4%pfa:CB+SBH-4%pfa:CB | -14.123007 | -27.576062 | -0.669951 | 0.0308279 | Yes |
| 4%pfa:Ctrl-4%pfa:CB | -25.944967 | -39.398023 | -12.491912 | 0.0000012 | Yes |
| 4%pfa:SBH-4%pfa:CB | -36.268105 | -49.72116 | -22.815049 | 0 | Yes |
| 4%pfa:Ctrl-4%pfa:CB+SBH | -11.821961 | -25.275016 | 1.6310948 | 0.1459944 | No |
| 4%pfa:SBH-4%pfa:CB+SBH | -22.145098 | -35.598154 | -8.6920425 | 0.0000359 | Yes |
| 4%pfa:SBH-4%pfa:Ctrl | -10.323137 | -23.776193 | 3.1299183 | 0.329417 | No |
| 10%f:CB+SBH-10%f:CB | 4.25424837 | -9.1988072 | 17.7073039 | 0.9994179 | No |
| 10%f:Ctrl-10%f:CB | -22.837124 | -36.29018 | -9.3840686 | 0.0000194 | Yes |
| 10%f:SBH-10%f:CB | -37.804444 | -51.2575 | -24.351389 | 0 | Yes |
| 10%f:Ctrl-10%f:CB+SBH | -27.091373 | -40.544428 | -13.638317 | 0.0000004 | Yes |
| 10%f:SBH-10%f:CB+SBH | -42.058693 | -55.511748 | -28.605637 | 0 | Yes |
| 10%f:SBH-10%f:Ctrl | -14.96732 | -28.420376 | -1.5142647 | 0.0162849 | Yes |
| 20%f:CB+SBH-20%f:CB | -7.1879739 | -20.641029 | 6.2650817 | 0.8734535 | No |
| 20%f:Ctrl-20%f:CB | -27.435556 | -40.888611 | -13.9825 | 0.0000003 | Yes |
| 20%f:SBH-20%f:CB | -43.62841 | -57.081465 | -30.175354 | 0 | Yes |
| 20%f:Ctrl-20%f:CB+SBH | -20.247582 | -33.700637 | -6.7945261 | 0.0001931 | Yes |
| 20%f:SBH-20%f:CB+SBH | -36.440436 | -49.893491 | -22.98738 | 0 | Yes |
| 20%f:SBH-20%f:Ctrl | -16.192854 | -29.64591 | -2.7397985 | 0.0061561 | Yes |
| 95%EtOH:CB+SBH-95%EtOH:CB | 9.34501089 | -4.1080447 | 22.7980664 | 0.5008772 | No |
| 95%EtOH:Ctrl-95%EtOH:CB | -9.295817 | -22.748873 | 4.1572386 | 0.5102015 | No |
| 95%EtOH:SBH-95%EtOH:CB | 1.57494554 | -11.87811 | 15.0280011 | 1 | No |
| 95%EtOH:Ctrl-95%EtOH:CB+SBH | -18.640828 | -32.093883 | -5.1877723 | 0.0007856 | Yes |
| 95%EtOH:SBH-95%EtOH:CB+SBH | -7.7700654 | -21.223121 | 5.6829902 | 0.7908445 | No |
| 95%EtOH:SBH-95%EtOH:Ctrl | 10.8707625 | -2.582293 | 24.3238181 | 0.2501752 | No |
| EMS:CB+SBH-EMS:CB | -39.000087 | -52.453143 | -25.547032 | 0 | Yes |
| EMS:Ctrl-EMS:CB | -58.055425 | -71.50848 | -44.602369 | 0 | Yes |
| EMS:SBH-EMS:CB | -72.748192 | -86.201247 | -59.295136 | 0 | Yes |
| EMS:Ctrl-EMS:CB+SBH | -19.055338 | -32.508393 | -5.6022821 | 0.0005486 | Yes |
| EMS:SBH-EMS:CB+SBH | -33.748105 | -47.20116 | -20.295049 | 0 | Yes |
| EMS:SBH-EMS:Ctrl | -14.692767 | -28.145822 | -1.2397113 | 0.0201049 | Yes |

**Supplemental Table 3:** Relevant Tukey's HSD statistical details on red autofluorescence ~ fixative*quenching. $\alpha$=0.05.

| Pairwise Comparison | Mean Difference | Lower Hinge | Upper Hinge | Adjusted P Value | Significant? |
|---|---|---|---|---|---|
| 4%pfa:CB+SBH-4%pfa:CB | -7.1691503 | -16.31833 | 1.98002953 | 0.2961427 | No |
| 4%pfa:Ctrl-4%pfa:CB | -1.1801743 | -10.329354 | 7.96900556 | 1 | No |
| 4%pfa:SBH-4%pfa:CB | -12.260915 | -21.410095 | -3.1117352 | 0.0013291 | Yes |
| 4%pfa:Ctrl-4%pfa:CB+SBH | 5.98897603 | -3.1602038 | 15.1381559 | 0.6042016 | No |
| 4%pfa:SBH-4%pfa:CB+SBH | -5.0917647 | -14.240945 | 4.05741515 | 0.8336959 | No |
| 4%pfa:SBH-4%pfa:Ctrl | -11.080741 | -20.229921 | -1.9315609 | 0.0056729 | Yes |
| 10%f:CB+SBH-10%f:CB | 6.78867102 | -2.3605088 | 15.9378509 | 0.3854476 | Yes |
| 10%f:Ctrl-10%f:CB | -10.891721 | -20.040901 | -1.7425413 | 0.0071086 | Yes |
| 10%f:SBH-10%f:CB | -16.420741 | -25.569921 | -7.2715609 | 0.000006 | Yes |
| 10%f:Ctrl-10%f:CB+SBH | -17.680392 | -26.829572 | -8.5312123 | 0.0000012 | Yes |
| 10%f:SBH-10%f:CB+SBH | -23.209412 | -32.358592 | -14.060232 | 0 | Yes |
| 10%f:SBH-10%f:Ctrl | -5.5290196 | -14.6782 | 3.62016025 | 0.7302007 | No |
| 20%f:CB+SBH-20%f:CB | -13.19207 | -22.34125 | -4.0428899 | 0.0004071 | Yes |
| 20%f:Ctrl-20%f:CB | -19.887582 | -29.036762 | -10.738402 | 0.0000001 | Yes |
| 20%f:SBH-20%f:CB | -29.337255 | -38.486435 | -20.188075 | 0 | Yes |
| 20%f:Ctrl-20%f:CB+SBH | -6.695512 | -15.844692 | 2.45366787 | 0.4092248 | No |
| 20%f:SBH-20%f:CB+SBH | -16.145185 | -25.294365 | -6.9960053 | 0.0000086 | Yes |
| 20%f:SBH-20%f:Ctrl | -9.4496732 | -18.598853 | -0.3004934 | 0.0364347 | Yes |
| 95%EtOH:CB+SBH-95%EtOH:CB | 1.83389978 | -7.3152801 | 10.9830796 | 0.9999993 | No |
| 95%EtOH:Ctrl-95%EtOH:CB | -4.1815686 | -13.330749 | 4.96761122 | 0.9638366 | No |
| 95%EtOH:SBH-95%EtOH:CB | 3.0908061 | -6.0583738 | 12.239986 | 0.9986403 | No |
| 95%EtOH:Ctrl-95%EtOH:CB+SBH | -6.0154684 | -15.164648 | 3.13371144 | 0.5967071 | No |
| 95%EtOH:SBH-95%EtOH:CB+SBH | 1.25690632 | -7.8922735 | 10.4060862 | 1 | No |
| 95%EtOH:SBH-95%EtOH:Ctrl | 7.27237473 | -1.8768051 | 16.4215546 | 0.2743122 | No |
| EMS:CB+SBH-EMS:CB | -42.988976 | -52.138156 | -33.839796 | 0 | Yes |
| EMS:Ctrl-EMS:CB | -38.079434 | -47.228613 | -28.930254 | 0 | Yes |
| EMS:SBH-EMS:CB | -51.946318 | -61.095498 | -42.797138 | 0 | Yes |
| EMS:Ctrl-EMS:CB+SBH | 4.90954248 | -4.2396374 | 14.0587223 | 0.8696027 | No |
| EMS:SBH-EMS:CB+SBH | -8.957342 | -18.106522 | 0.19183781 | 0.0608723 | No |
| EMS:SBH-EMS:Ctrl | -13.866885 | -23.016064 | -4.7177047 | 0.0001703 | Yes |

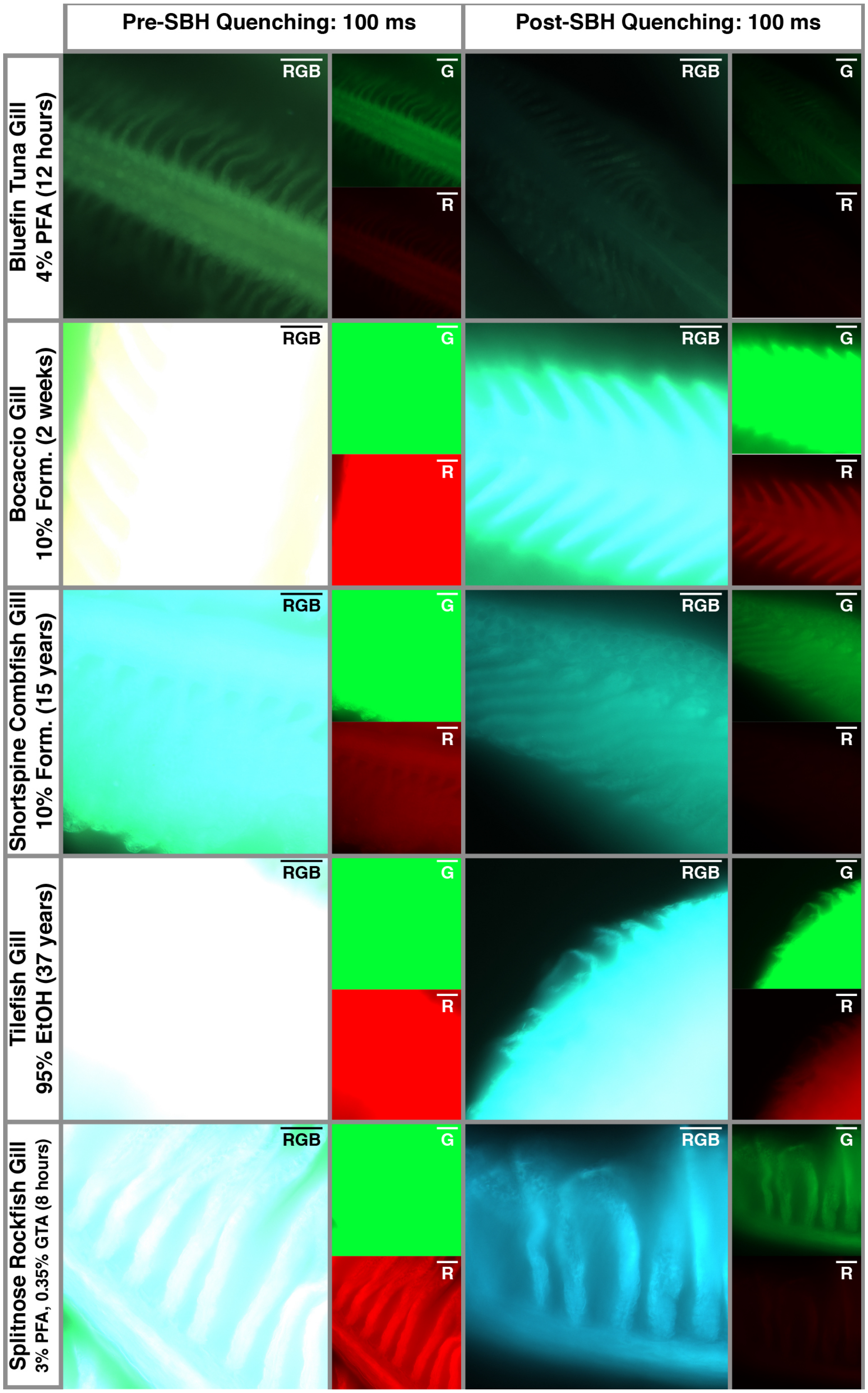


**Supplemental Figure 1:** Whole-mount epifluorescence images of various archival fish gills before and after sodium borohydride (SBH) quenching.

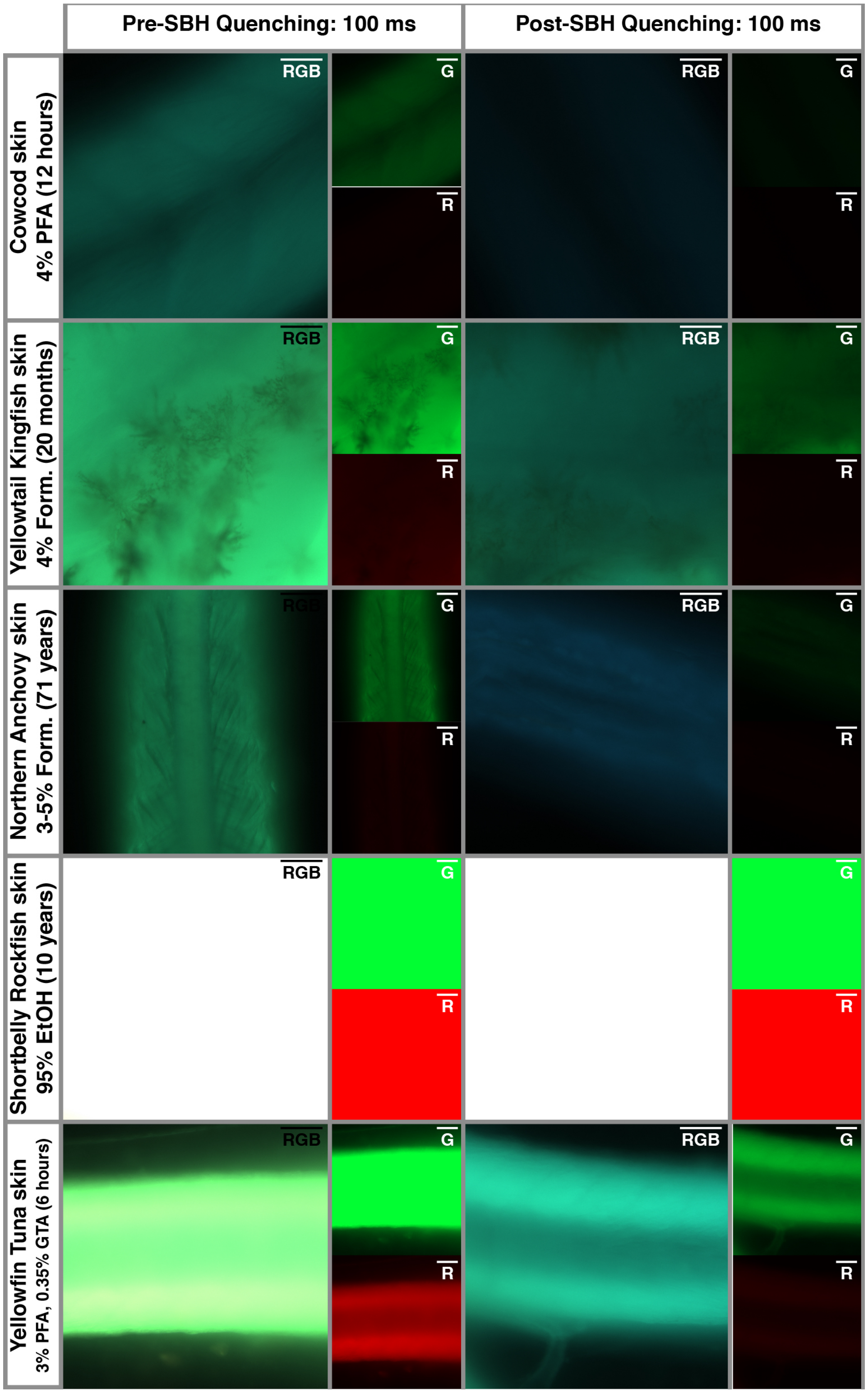


**Supplemental Figure 2:** Whole-mount epifluorescence images of various archival fish skin before and after sodium borohydride (SBH) quenching.
